# Challenges and advances for transcriptome assembly in non-model species

**DOI:** 10.1101/084145

**Authors:** Arnaud Ungaro, Nicolas Pech, Jean-François Martin, R.J. Scott McCairns, Jean-Philippe Mévy, Rémi Chappaz, André Gilles

## Abstract

Analyses of high-throughput transcriptome sequences of non-model organisms are based on two main approaches: de *novo* assembly and genome-guided assembly using mapping to assign reads prior to assembly. Given the limits of mapping reads to a reference when it is highly divergent, as is frequently the case for non-model species, we evaluate whether using blastn would outperform mapping methods for read assignment in such situations (>15% divergence). We demonstrate its high performance by using simulated reads of lengths corresponding to those generated by the most common sequencing platforms, and over a realistic range of genetic divergence (0% to 30% divergence). Here we focus on gene identification and not on resolving the whole set of transcripts (i.e. the complete transcriptome). For simulated datasets, the transcriptome-guided assembly based on blastn recovers 94.8% of genes irrespective of read length at 0% divergence; however, assignment rate of reads is negatively correlated with both increasing divergence level and reducing read lengths. Nevertheless, we still observe 92.6% of recovered genes at 30% divergence irrespective of read length. This analysis also produces a categorization of genes relative to their assignment, and suggests guidelines for data processing prior to analyses of comparative transcriptomics and gene expression to minimize potential inferential bias associated with incorrect transcript assignment. We also compare the performances of *de novo* assembly alone *vs* in combination with a transcriptome-guided assembly based on blastn via simulation and empirically, using data from a cyprinid fish species and from an oak species. For any simulated scenario, the transcriptome-guided assembly using blastn outperforms the *de novo* approach alone, including when the divergence level is beyond the reach of mapping methods. Combining *de novo* assembly and a related reference transcriptome for read assignment also addresses the bias/error in contigs caused by the dependence on a related reference alone. Empirical data corroborate those findings when assembling transcriptomes from the two non-model organisms: *Parachondrostoma toxostoma* (fish) and *Quercus pubescens* (plant). For the fish species, out of the 31,944 genes known from *D. rerio*, the guided and *de novo* assemblies recover respectively 20,605 and 20,032 genes but the performance of the guided assembly approach is much higher for both the contiguity and completeness metrics. For the oak, out of the 29,971 genes known from *Vitis vinifera*, the transcriptome-guided and *de novo* assemblies display similar performance but the new guided approach detects 16,326 genes where the *de novo* assembly only detects 9,385 genes.

## Introduction

Synthesis and maturation of RNAs is an elemental cog in the cellular machinery. Although inherently noisy, transcriptional variation can be associated with basal/fundamental processes such as enzyme activity [1] and protein production [2]. Transcriptional variation may also underlie complex morphological differences [3,4], and may itself be a target for natural selection [5]. As such, the quantification of RNA abundance remains an essential link in deciphering the genotype-phenotype map. In this context, transcriptome inference (i.e. *in silico* assembly and annotation) is an initial and requisite basis for studying gene expression [6,7]. After two decades of RNA microarrays [8], RNA-seq has democratized the analysis of transcriptomes for any non-model organism. This technological innovation has spread to several new uses in multiple domains in the life sciences, from direct applications such as transcript annotation [9,10], to providing insights into *cis* and *trans* regulation in allopolyploid species [11], speciation [11,12], heat stress [13], ecotoxicology [14] and ecology and evolution in general [15].

Although RNA-seq may be applied to non-model organisms, meaningful transcriptome inference in the absence of a reference genome is not a trivial problem. The two most common approaches are *de novo* assembly (i.e. assembling reads free of any reference genome/transcriptome) and genome-guided transcriptome assembly (i.e. mapping reads to a related reference genome to identify transcript models, then assembling those transcripts). Cahais *et al*.[16] reconstructed the transcriptomes of non-model organisms through *de novo* assembly, combining 454 and Illumina sequence reads, and explored the efficiency of the approach through annotating the assembled contigs against the taxonomically closest reference genome. Although they retrieved a great number of transcripts, and opened new opportunities of transcriptome inference for non-model organisms, they also found potential issues, namely: sensitivity of alignment error due to paralogs and multigene families; production of artefactual chimeras; problems reconstructing transcript length, and potentially misestimating allelic diversity. These issues were further confirmed in several subsequent articles dealing with the *de novo* strategy [17–19], although the extent of sensibility to each varies slightly depending on the software and analytical solutions used.

Vijay *et al*. [20] compared *de novo* and genome-guided transcriptome assemblies, concluding that the genome-guided approach (based on mapping reads to a reference genome prior to assembly) is less sensitive to sequencing error rate and polymorphisms. It is also less sensitive to paralogous loci than *de novo*. However, by simulating polymorphic sequences, they also showed that the accuracy of guided transcriptome assemblies is highly sensitive to genetic divergence between the reads and the reference genome, with significant declines in the performance of mapping when sequence divergence exceeds 15% [20]. Likewise it was also demonstrated that evolved structural differences between reference and query (e.g. indels, inversions) can generate bias/error in transcript sequences inferred via guided assembly, due to their dependence on the reference as a template in mapping [20,21]. Finally, Jain *et al*. [22] proposed a combination of *de novo* and genome-guided approaches wherein transcripts from Trinity (*de novo*) were augmented with those from TopHat1-Cufflinks (genome-guided). Although this approach increased the overall efficiency of transcriptome inference, it did not address the specific limitations of each method. The current challenge is therefore inferring transcriptomes while addressing those limitations, in particular when divergence with the closest related genome (or transcriptome) is high, as is typically the case for non-model organisms. Guided assembly has considerable potential [23], but the effect of divergence on mapping efficiency may be a critical factor limiting this potential. We decided to address this issue by combining the *de novo* approach with a modified guided transcriptome step. We chose an assignment method that would be less sensitive to divergence, namely the widely used nucleotide BLAST (blastn) algorithm. Blastn finds regions of similarity between biological sequences and can accept more relaxed similarities than mapping procedures. In theory, this would make it a tool of choice in assigning reads to genes prior to assembly, and could overcome the limitations of mapping in the guiding step.

Here we first test for the performance of a read assignment pipeline based on blastn, using simulated reads generated from a well-characterized reference transcriptome. We estimate the assignment error rate of a read by simulating a range of read lengths corresponding to those generated by the most common sequencing platforms (100, 150, 200, 350 bases), and over a realistic range of genetic divergence between the simulated reads and the reference transcriptome (0%, 5%, 15% and 30%). Whatever the approach used, genomic complexity can create difficulties in reconstructing transcriptomes. In particular, genome duplications exacerbate biases in transcriptome assembly because of the higher number of paralogous loci. Teleostean fishes in general, and cyprinids in particular, are good models for studying the effects of paralogs on transcriptome assembly as they display a complex genome with multiple rounds of duplication [24]. We therefore chose *Danio rerio* (Teleostean: Cyprinidae) as a reference transcriptome for this *in silico* analysis. Once we characterized the performance of blastn in assigning reads, we compare the performances of *de novo* transcriptome inference alone and in combination with blastn as a method to assign reads to genes prior to assembly. This is done first on the same simulated data as previously described, using *Danio rerio* as a reference and then applied to two non-model organisms to ensure generality. We first applied the method to a cyprinid fish species, *Parachondrostoma toxostoma* (Vallot, 1837), keeping *Danio rerio* as the reference transcriptome for read assignment, as it was demonstrated that there are large synteny blocks among Cyprinidae fish [25]. We further tested the method on a plant species, *Quercus pubescens*, using *Vitis vinifera* as a reference transcriptome [26]. This distant reference transcriptome was selected to facilitate comparison with Torre et al. analysis [26]. This allows for an empirical comparison of the relative performance of the two methods in the face of high divergence between an inferred transcriptome and a related reference transcriptome for a broad spectrum of organisms. This whole analysis addresses whether the transcriptome-guided assembly using blastn improves transcriptome inference for non-model organisms.

## Materials and Methods

### Implementing read assignment with blastn

We implemented a flexible assignment pipeline (Fig 1) as follows; note that all scripts are available at https://github.com/egeeamu/voskhod. Sequencing reads are filtered for over-representation (PCR duplicates or over-expression) using a custom Python script and then merged, when relevant (i.e. when using overlapping paired-end reads), with Pear [27] using the following parameters: minimum overlap -v 8, scoring method -s 2, p-value -p 0.01, minimum assembly length -n 30, maximum assembly length -m 0. After merging – a step in the pipeline that is optional – all available reads are quality filtered using a custom script that first trims 5’ and 3’ ends on the basis of Phred quality score, trimming bases until a Phred score is ≥ 13 (i.e. < 0.05 error rate) is encountered. A read is further rejected (and replaced by a sequence made up by 50 “Ns” to keep the parity between R1 and R2 files when it applies) if it displays one of the following conditions:

i. one base with a Phred quality score < 5;
ii. more than 10% of bases with a Phred quality score < 13;
iii. a mean Phred quality score < 20 for the entire read;
iv. a length shorter than 30 bases.

**Fig 1.**
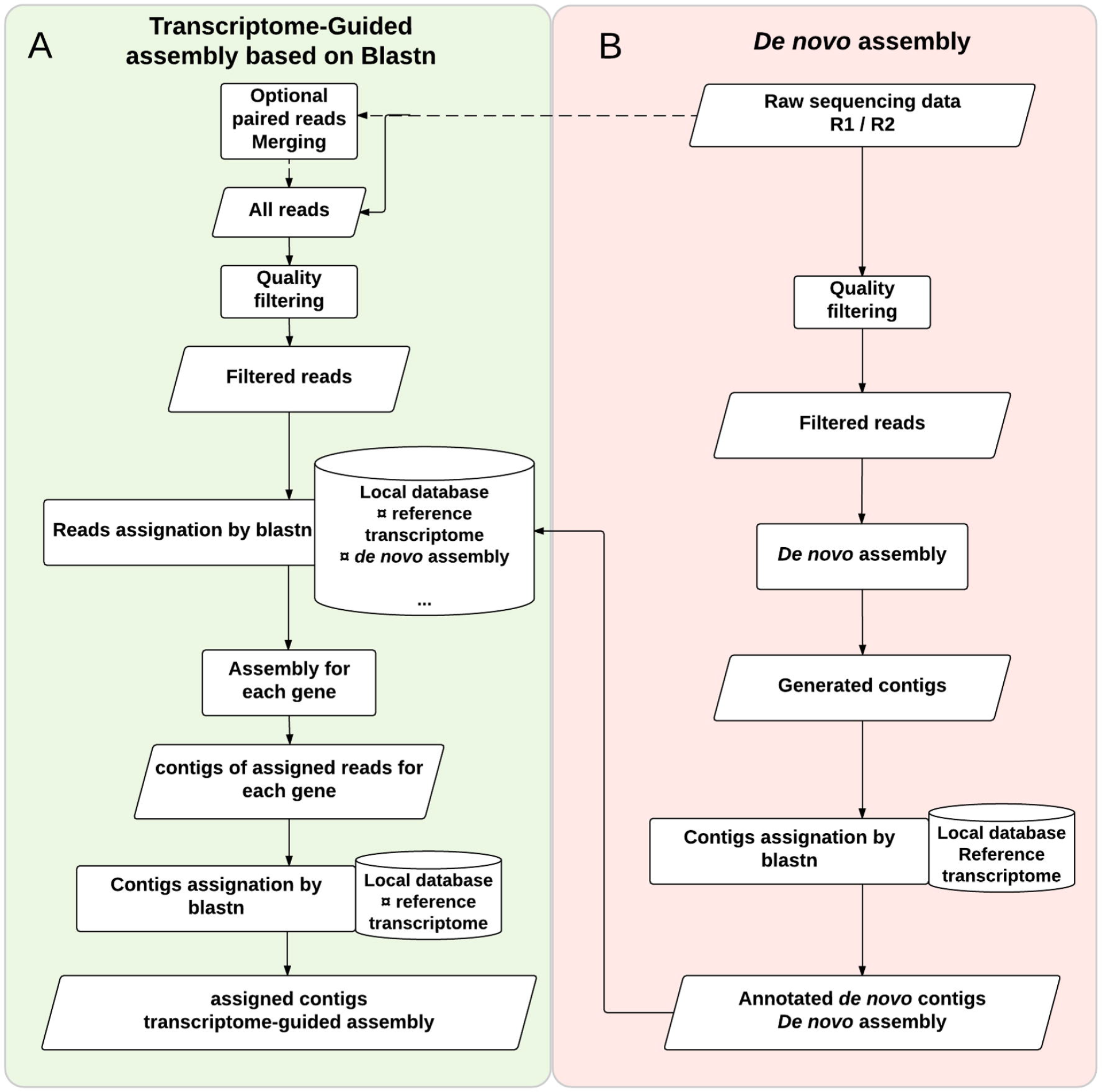
Transcriptome-guided assembly Pipeline. We present the pipeline for transcriptome assembly and contig assignment. (A) The transcriptome-guided assembly *per se* (green background) combines a **read assignment step** based on blastn, making use of a local database merging the related reference transcriptome and *de novo* assembly of the query transcriptome. (B) The *de novo* assembly (red background) with a blastn annotation of generated contigs to obtain a *de novo* assembly for the inferred transcriptome. Cylinders represent the database, rectangles represent analytical steps in the processes, and parallelograms correspond to results. Note that the dashed lines represent optional steps in the pipelines.

Next, Nucleotide-Nucleotide BLAST [28] is used to assign filtered reads to *gene-id(s)* by finding regions of similarity between the read and the reference transcriptome(s) stored in a local SQLite database [29] with Word-size = 9, HSP > 70% and similarity > 70%. This set of parameters allows for filtering chimeras and contaminants when applied to reference transcriptomes. With such parameters, it would be inadequate to use a genome as reference because individual reads could match multiple exons from the same gene while being erroneously rejected from the analysis. In the following procedures, we therefore use reference transcriptomes only. Of course, when an annotated genome is available, one can extract transcripts to reconstruct the corresponding reference transcriptome. Each read is assigned to the *gene-id* corresponding to the best hit. When identical scores (overall quality of an alignment) are obtained for multiple *gene-ids*, this information is stored in the variable *Hit_multigene*. This variable is used to reflect the level of mismatches on paralogous genes or multigene families. As the pipeline is designed to accommodate multiple reference transcriptomes, when identical scores are obtained for a single *gene-id* from distinct reference transcriptomes in the local database, this information is stored in the variable *Hit_multispecies* along with the associated *gene-id*. This variable is used to reflect the level of mismatches on orthologous genes when relevant. The final output is a collection of database entries displaying the number of assigned reads for each *gene-id* (see supporting information 1 for an example) stored in the local database and exported in a tabular file for further analysis.

### Measuring blastn efficiency and performance for read assignment

Based on the published transcriptome of *Danio rerio* (extracted from genome version Zv10), we used the longest transcript for each gene (i.e. 31,953 transcripts) [30] and simulated RNA-seq reads of known variability. We simulated reads for four length classes (100, 150, 200 and 350 bases, see supporting information 2 for details), with each dataset corresponding to the whole transcriptome at 10X coverage for each gene. Varying read lengths allows studying erroneous assignments caused by conserved homologous regions (e.g. causing assignment to multiple values for *gene-id*), the expected outcome being that longer reads will decrease ambiguity. Sequence divergence between simulated reads and the reference transcriptome was used as a proxy to simulate confounding processes such as sequence error rate (estimated at 0.64% for R1 and 1.07% for R2 for the Illumina Miseq sequencer; [31]), polymorphism (ranging from 0.1% to 1%; [20]), and species divergence. We mimicked sequence divergence by introducing random base errors into simulated reads at rates of 0%, 5%, 15% and 30%, as in Vijay *et al*. [20] (see supporting information 2 for details).

The efficiency (percentage of output to input) was estimated from the number of genes recovered with the read assignment pipeline (output), relative to the number of genes within the reference transcriptome (input). We assessed the performance (i.e. quality of the output) with two metrics:

i. the recovery rate (*rr*), defined as the proportion of reads simulated from a given *gene-id* and correctly assigned to this *gene-id*
ii. the specificity rate (*sr*), defined as the number of reads assigned to a *gene-id* that were simulated from this *gene-id* relative to the total number of reads assigned to that *gene-id*

For example, consider 100 reads simulated from a given *gene-id*. If the pipeline assigns 80 from these reads to the same *gene-id*, then the *rr* is estimated as 80/100 = 0.8. Conversely, if we observe 150 reads assigned to this *gene-id* (80 reads from the same *gene-id* as source and 70 reads from other *gene-id(s*)) the specificity rate *sr* is estimated as 80/150 = 0.53.

We propose a typology of genes based on the assignment of reads considering their recovery rate (*rr*) and specificity rate (*sr*).

i. The category *rr*=1 and *sr*=1 constitutes the ‘perfect’ genes, all generated reads for a gene are recovered for this gene and no read from another gene is “captured”
ii. The category *rr*=1 and *sr*<1 constitutes the ‘recipient’ genes, all the generated reads for a gene are recovered for this gene and at least one read from another gene is “captured”
iii. The category *rr*<1 and *sr*=1 constitutes the ‘donor’ genes, at least one generated read for a gene is not recovered for this gene and no read from another gene is “captured”
iv. The category *rr*<1 and *sr*<1 constitutes the ‘mixed’ genes, at least one generated read for a gene is not recovered for this gene and at least one read from another gene is “captured”
v. The category *rr*=0 and *sr*=N.A. constitutes the ‘undetectable’ gene, no assigned reads for this gene although they were generated

Finally, as gene length is highly heterogeneous in transcriptomes [32], we tested whether assignment success is impacted by this factor, with each gene’s length estimated as the length of its longest transcript.

The aforementioned performance metrics (*rr* & *sr*) were modeled as a function of gene length (treated as a continuous variable), read length, sequence divergence (each treated as categorical variables) and all interaction terms amongst factors using a mixed-effect logistic regression implemented in the lme4 package [33] for R (version 3.3.1; [34]). Aforementioned model terms were treated as fixed-effects, and gene identity was included as a random factor (see supporting information 3 for details).

### Transcriptome assembly approaches

#### De novo assembly pipeline

In a recent study, Lu et al. [35] compared various assembly software, including: Trinityrnaseq_r2012-04-27 [9], Oases [36] and trans-ABySS [37]. The authors demonstrated that Trinity produces assemblies with the highest completeness and contiguity (using default parameters). Wang and Gribskov [38] also rank Trinity among the *de novo* assembly programs producing fewer chimeras. Our *de novo* assembly (Fig 1B) is built therefore on Trinity [9,39]. Both single-end and paired-end reads can be used. When paired-end reads are used, merging the R1 and R2 reads is unnecessary as Trinity uses the information from separate R1 and R2 paired files. The quality-based filtering procedure was the same as for the read assignment pipeline, as implemented specifically for non-merged reads. The *de novo* assembly was done using Trinity 2.2.0 with *normalize_reads* and *stranded_library* options, using the combination of R1 and R2 reads when available. The output of the assembly is a collection of generated contigs stored in the local database and exported in a single FASTA file.

Each contig was first identified/annotated by alignment against the transcriptome of the reference species using blastn (Word-size = 9, HSP > 70% and similarity > 70%), and used to populate a reference database. Only the best HSP was conserved and converted to reverse-complement when a strand differed from the reference. When multiple hits occurred (i.e. the same blastn score), the contig was assigned to the corresponding *gene-ids* and denoted with the variable *Hit_multigene* in the database. This step is crucial to remove gap effects in some paralogous sequences with conserved regions. The output of this step is a collection of assigned contigs stored in the local database (i.e. the reconstructed *de novo* assembly) and exported as FASTA files, one for each *gene-id*. These files define a *de novo* inferred transcriptome of the non-model species that was stored in the local database.

#### Developing a transcriptome-guided assembly pipeline based on blastn

Given the impact of divergence between the reference transcriptome and sequencing reads on assignment, and “because the resulting assembly can be biased towards the closely related genome rather than the focal genome” [21] (<u>i.e. constraining the annotation to the reference structure of transcripts</u>), it is interesting to include as reference the collection of annotated contigs issued from a *de novo* assembly produced with Trinity, in combination with the related reference transcriptome. With this design, the assignment profits both from sequence proximity with the *de novo* contigs, thereby decreasing the effect of sequence divergence, while simultaneously allowing for assigning reads to genes not reconstructed *de novo*, hence increasing transcriptome coverage.

First, the *de novo* approach was used as previously described, producing a list of assigned contigs in a local database that also contains the related reference transcriptome (Fig 1B). The reads, merged with Pear when working with paired end sequencing, are filtered on quality and assigned to the local reference database. Each read (merged or not) is assigned with blastn to the *gene-id* corresponding to its best hit as described in the read assignment pipeline. Each assigned read is stored in the local database and appended to a FASTA file (one file per gene) before assembly. To avoid circularity with the *de novo* assembly, we used successively Spades (v3.7.1.8; *careful* option and *cov-cutoff* = auto; [40] with the auxiliary BayesHammer error correction algorithm [41], and then CAP3 [42] default parameters). We used a combination of these two established software solutions because they rely on different assumptions with different behaviors regarding read length, quantity of reads and transcript diversity [16,35,43,44]. The output of this step (one SQLite file; output from each assembly program concatenated into a single file) is a collection of contigs. A final validation step was used to prevent chimeric or mis-assembled reconstructions generated during the assembly step. All contigs were annotated using blastn and the related reference transcriptome stored in the local database. Contigs not annotated at this step were discarded (in practice they mostly involved repetitive domains). Retained contigs and HSPs were conserved and reverse-complemented to fit the reference orientation when necessary. The final output is a collection of annotated contigs stored in the local database as well as FASTA files (one for each *gene-id*).

### Assessing the quality of inferred assemblies

In this section, we compare the performance of *de novo* assembly alone (Fig 1B) and the transcriptome-guided assembly pipeline based on blastn (Fig 1A). This comparison is made first on assemblies based on reads simulated from the *D. rerio* transcriptome, and then on empirical data from the two non-model species (*Paratoxostoma toxostoma*, a cyprinid fish species and *Quercus pubescens*, an oak species). We tested the impact of sequence divergence both over a range of simulated values (0%, 5%, 15% and 30%), and also using extant inter-specific divergence between the reference transcriptome and the analyzed species, respectively *D. rerio* for *P. toxostoma* and *V. vinifera* for *Q. pubescens*. We used fixed read lengths of 100 and 200 nucleotides at a homogeneous coverage of 10X for simulated data; when for empirical data, median read length was 234 bases for *P. toxostoma* and 192 bases for *Q. pubescens*, while depth of coverage was obviously variable.

A diversity of metrics should be used when assessing the quality of inferred transcriptomes. Although these metrics have not been definitively established, accuracy, completeness, contiguity, chimerism and variant resolution capture all essential elements of transcriptome quality [45]. These metrics are implemented in Rnnotator [46], and were used in the comparative study of *de novo* and genome-guided assembly by Lu et al. [35]. We have adapted some of these metrics for assessing transcriptome assemblies as we focus on genes and not on transcripts. We define:

i. The number of identified genes (*NIG*). A gene is considered as identified if at least one contig is assigned to this gene.
ii. The Completeness (Cp) for a gene *g* corresponds to the proportion of the length for the longest reference transcript (*L_g_*, in number of bases) covered by the whole set of aligned contigs for the gene *g* (*C_gj_*, *j*=1…*n_g_*). The completeness is maximal (equal to 1) when the combined length of the aligned contigs (using ProbCons, [47]) matches the longest reference transcript.

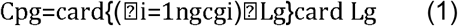
iii. The Contiguity (Ct_g_) for a gene *g* corresponds to the proportion of the longest reference transcript (*L_g_*, in number of bases) covered by the longest contig for this gene (*C_gjmax_*). The contiguity is maximal (equal to 1) when there is a perfect match between the longest assigned contig and the longest reference transcript.

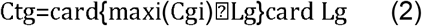

These last two metrics are adapted from Martin & Wang [45] because we focus on gene identification for non-model organisms and not on resolving the whole set of transcripts (i.e. the complete transcriptome). We used the longest reference transcript for each gene in the reference transcriptome (*D. rerio* or *V. vinifera*) as an upper bound of transcript length for the contigs from RNA-seq. Variation in contiguity and completeness was analyzed using a linear mixed-effect model with divergence, assembly approach (*de novo* assembly or transcriptome-guided assembly using blastn) and their interaction treated as fixed factors, and random variation attributed to gene. This was done both for simulated and empirical data. For simulated data, we modeled the four divergence levels (0%, 5%, 15%, 30%) as fixed factors. For empirical data, we characterized the distribution of the divergence between each species and its reference transcriptome for each gene and did not model it explicitly. Finally, biological processes associated with each gene was inferred based on annotations from the Protein ANalysis THrough Evolutionary Relationships database (http://pantherdb.org/about.jsp; [48,49]) and are provided as supporting information 9.

### Biological material & empirical data

All sampling and experimental protocols were reviewed and approved by local regulatory agencies (ONEMA and the DDT from Alpes-de-Haute-Provence, Hautes-Alpes and Vaucluse; authorization number 2007–573 and 2008–636, following national regulations. Samples of *Parachondrostoma toxostoma* (3 males) were collected from two rivers: the Durance (southern France) and Ain (eastern France). Eight tissues were sampled (liver, hindgut, midgut, heart, brain, gill, caudal fin, spleen) from one specimen (euthanized by decapitation), only the caudal fin was sampled from the two other males (non-invasive sampling). Samples were ground in liquid nitrogen and total cellular RNA was extracted using the RNeasy Plus Universal kit (Qiagen). The TruSeq Stranded mRNA Library Preparation kit (Illumina Inc., USA) was used according to the manufacturer’s protocol, with a few modifications (see supporting information 4 for details). Samples were uniquely barcoded for individuals and tissues, and pooled cDNA libraries were sequenced using a Miseq Illumina sequencer with the 2 x 250 paired-end cycle protocol (see supporting information 4 for details). In total, 16,216,379 reads were used to reconstruct the transcriptome of *P. toxostoma*. All data acquired for this study are available as an SRA archive at https://trace.ncbi.nlm.nih.gov/Traces/sra/sra.cgi?study=SRP091996 (SRX2266500 to SRX2266509).

For plant samples, we used five leaves belonging to a single *Quercus pubescens* specimen from the Oak Observatory at the Observatoire de Haute Provence (France). Total RNA was extracted using the RNeasy Plant mini Kit (Qiagen). The TruSeq Stranded mRNA Library Preparation kit (Illumina Inc., USA) was used according to the manufacturer’s protocol. Libraries were sequenced using a Next-Seq-500 Illumina sequencer with the 2 x 150 paired-end cycle protocol. In total, 46,881,297 reads were used to reconstruct the transcriptome of *Q. pubescens*. All data acquired for this study are available as an SRA archive at https://trace.ncbi.nlm.nih.gov/Traces/sra/sra.cgi?run=SRR5410765

## Results

### Evaluation of the read assignment pipeline with simulated data

The mixed logistic model describing recovery rate (*rr*) was highly significant (*X^2^*=27,605,272, df=23, P< 2.2e^-16^), with each term of the model being significant (supporting information 5). Recovery rate (*rr*) increased as a function of gene length irrespective of read length and divergence (Fig 2); however, the effect was most pronounced at 30% divergence between query and reference. At this divergence level, fewer than 50% of reads from genes smaller than 2kb (350 base reads; Fig 2G) to 4kb (100 bases; Fig 2A) – a substantial fraction of the transcriptome (Fig 3A) – were recovered. Likewise, high divergence was associated with the greatest variability in *rr* (dashed lines on Fig 2). Conversely, *rr* was generally greater than 0.9 under all other scenarios of divergence, including 15%. Read length influenced the shape of the curve describing the gene length at which *rr* approached unity, with longer reads yielding perfect scores for smaller genes (Figs 2G and 2H). As before, this effect was substantially reduced under the high divergence scenario.

**Fig 2.**
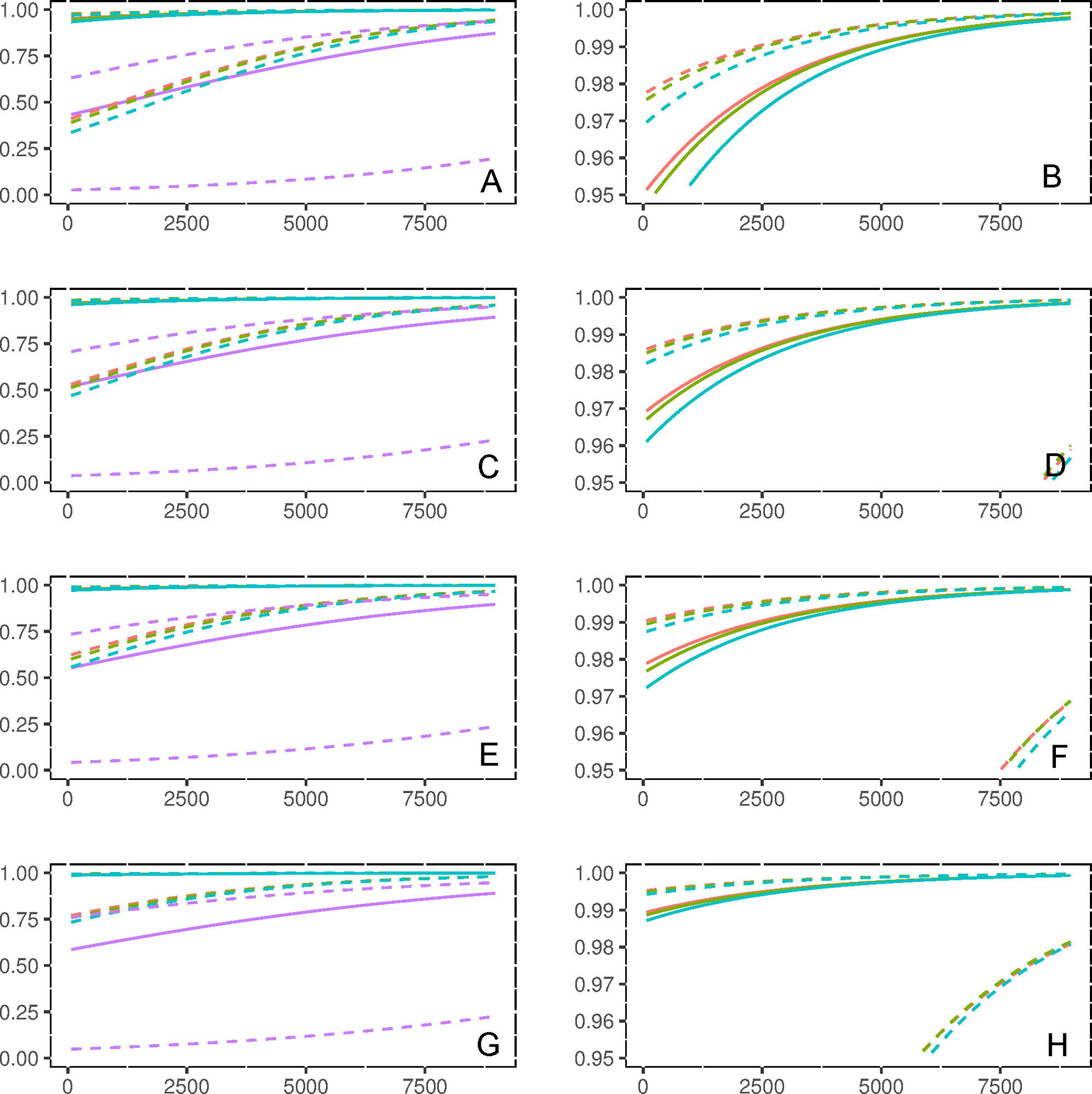
**Predicted recovery rate (rr)** using the mixed logistic modelization as a function of gene length (rr), read length and divergence between target and reference transcriptomes: red lines denote 0% divergence; green 5%; blue 15% & purple 30%. Solid lines correspond to the median of predictions, conditioned on random variation among genes, with 80% prediction intervals indicated by dashed lines. Read length increases downward and panels to the right represent a magnified view of the upper 5th quantile of rr scores to better visualize differences between low divergent sequences: 100 base reads (A & B); 150 bases (C & D); 200 bases (E & F); 350 bases (G & H).

**Fig 3.**
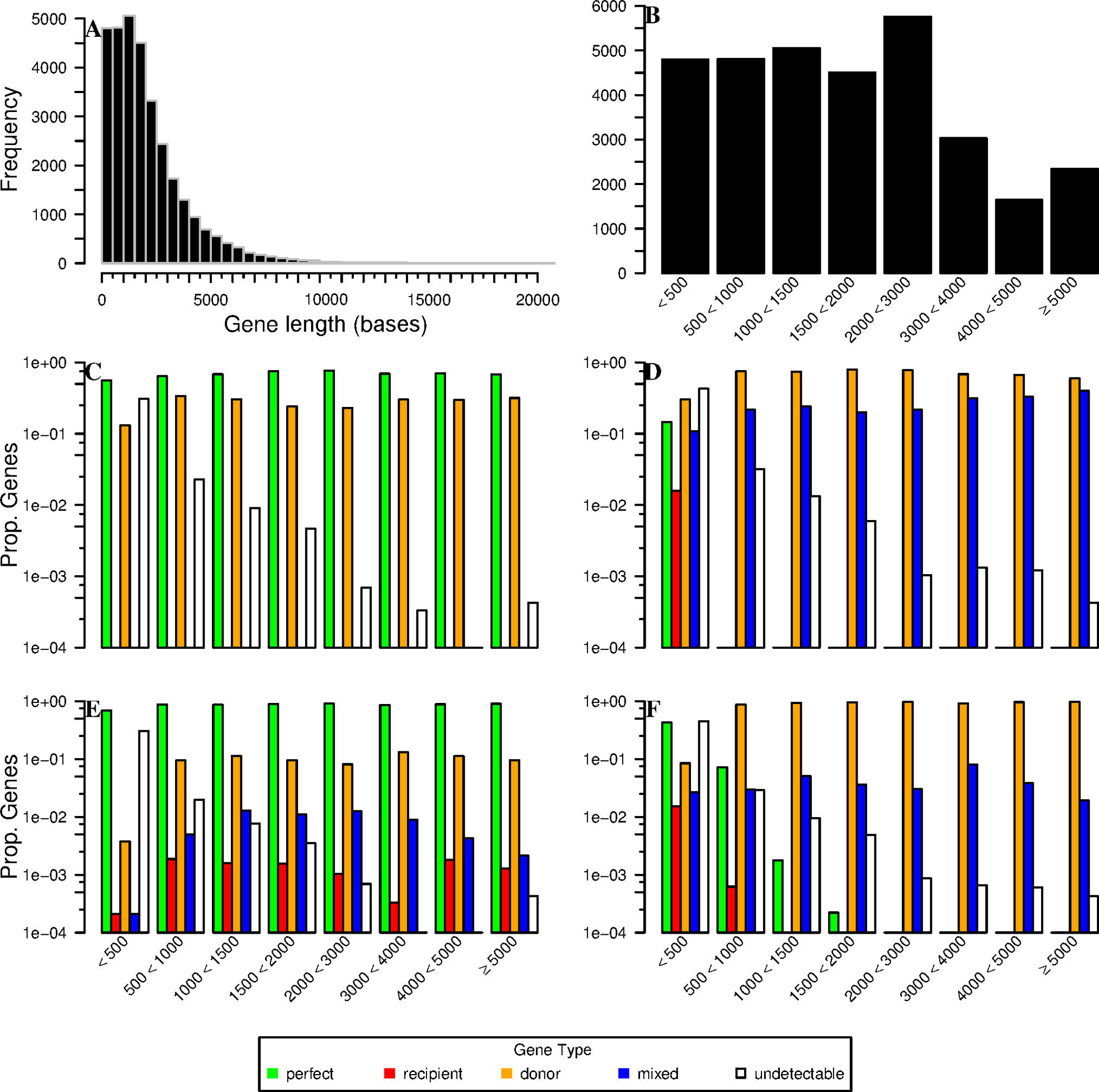
**Proportion of gene types recovered** in divergent simulations by size-class of gene. (A) Histogram of gene lengths for the *Danio rerio* transcriptome used for simulating RNA-seq reads. (B) Number of genes from A, by size-class in subsequent windows. 100 base reads are plotted with 0% divergence (C) and 30% divergence (D); 350 base reads with 0% divergence (E) and 30% divergence (F). Gene types are described in the legend.

The mixed logistic model for specificity rate (*sr*) was also highly significant (*X^2^*=1,625,516, df=8, P< 2.2e-16). Specificity rate displayed median values near to one (supporting information 5), with the random effect being greater (standard deviation estimate=4.00) compared to that for *rr* (standard deviation estimate=2.341). Each term of the model appeared significant (supporting information 5). Relations between factors and *sr* are the same as for *rr*, with greaterread and gene lengths being associated with higher values of *sr*, and lower *sr* under higher divergence.

The dataset of simulated reads used to evaluate read assignment performance consisted of 31,944 genes of varying length (Figs 3A and 3B). In general, the proportion of ‘perfect’ genes (*rr*=1, *sr*=1) increased with read length. For example, at 0% divergence between reference transcriptome and simulated reads 21,913 genes belonged to the ‘perfect’ category for read length of 100 bases (Fig 3C), whereas 27,289 were so classified for read lengths of 350 bases (*X^2^*=2554.4, df=1; P< 2.2e^-16^, Fig 3E). This improvement in performance coincided with a decrease in the proportion of ‘donor’ genes (*rr*<1, *sr*=1), but also with slight increases in the number of ‘recipient’ (*rr*=1, *sr*<1) and ‘mixed’ (*rr*<1, *sr*<1) gene classes (Fig 3E). Surprisingly, gene length also had a modest impact on improving assignment performance, with longer genes accumulating a slightly higher proportion of ‘perfect’ genes; however, this trend was only observed in the absence of divergence between reference transcriptome and simulated reads. Sequence divergence had a drastic impact on the proportion of genes that could be classed as perfect: at 30% divergence only 700 genes for reads of 100 bases (Fig 3D; 2.19% of all genes, *X^2^*=979.5, df=1; P< 2.2e^-16^) and 2,401 for read lengths of 350 bases (Fig 3F; 7.52%).

Increasing divergence also increased the ‘donor’ gene category (Figs 3D and 3F), detrimental to the ‘perfect’ gene category. Here increasing read length also appeared to slightly increase the proportion of ‘donor’ genes in general, with the maximum proportion of ‘donor’ genes observed for reads of 30% divergence and a length of 350 bases (81.04%, i.e. 25,888 genes); this also corresponded to a low proportion for the ‘perfect’ gene category (7.52%, i.e. 2,401 genes). The percentage of ‘mixed’ genes also increased with divergence, although this increase was most pronounced for short reads (Fig 3D) and was maximal for reads of 100 bases (7,370 genes; 23.07%).

Interestingly the proportion of ‘recipient’ genes (*rr*=1, *sr*<1) was low whatever the factor studied. The minimum value was 0% (no gene) at 0% divergence whatever the read lengths. The greatest number of ‘recipient’ genes (391 genes; 1.22%) was observed for reads of 15% divergence and a read length of 100 bases (supporting information 6). Likewise, the ‘undectable’ gene category (*rr*=0, *sr*=N.A.) displayed low proportions overall, ranging from 5.08% to 7.40% (i.e. 1,623 to 2,365 genes), with the greatest proportions being observed for short genes.

### Comparison between assembly approaches based on simulations

For reads of 100 bases, *de novo* assembly alone recovered a marginally higher number of genes (*NIG*) than the transcriptome-guided assembly pipeline based on blastn, irrespective of divergence with the reference transcriptome (Fig 4A): numbers for *de novo* ranged from 31,930 to 31,938 and from 31,147 to 31,944 for guided assembly. Conversely, measures of the quality of transcript assembly were higher for the guided assembly than for*de novo* alone. The completeness (*Cp*) of *de novo* assembled 100 base reads ranged from 0.74 to 0.89, with *Cp* generally increasing with sequence divergence (Fig 4B); *Cp* for guided assembly ranged from 0.97 to 0.99. Contiguity (*Ct*) exhibited similar patterns, increasing with divergence for *de novo* assemblies (Fig 4C; 0.67 to 0.87), yet was relatively stable, but higher for guided assembly (0.97 to 0.99). Increasing read length to 200 bases resulted in an overall decrease in the number of genes recovered, as well as a reversal in the patterns of pipeline performance, with guided assembly generally identifying more genes (28,339 to 28,532); *de novo* alone at the same levels of divergence tended to recover fewer genes (27,597 on average), with the exception of 28,604 genes at 15% divergence (Fig 4A). *Cp* and *Ct* were improved in both pipelines when read length was increased to 200 bases, although the guided assembly continually outperformed *de novo* alone at most levels of divergence (average *Cp* of 0.95 *vs* 0.99; average *Ct* of 0.89 *vs* 0.99), except at 30% where metrics were similar for each assembly approach.

**Fig 4.**
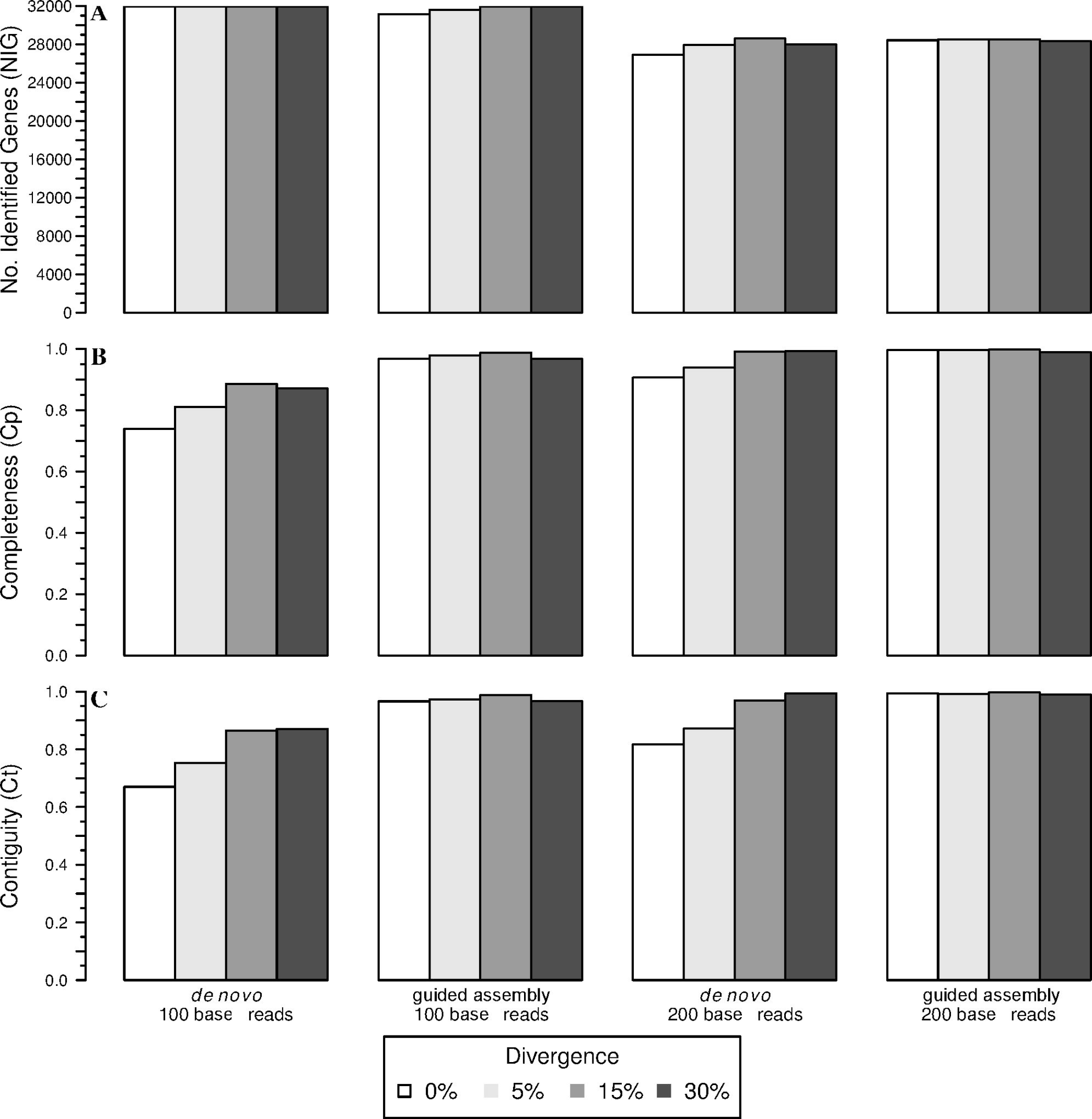
**Efficiency and performance of *de novo* and transcriptome-guided assembly based on Blastn** by read length and divergence level. (A) Number of identified genes; (B) completeness; (C) contiguity. Divergence between target and reference transcriptomes is described in the figure legend.

Fig 5 shows a non-parametric estimation of the bivariate distribution of contiguity and completeness for 0% divergence and read lengths 100 and 200 bases – note that for clarity, only ‘imperfect’ genes are displayed (the full range of factors is described in supporting information 7). For 100bp reads of 0% divergence, the *de novo* approach alone showed only 13,590 genes (46.1%) with a contiguity and a completeness equal to 1, while a large cluster of genes was observed with quality metrics inferior to 0.1 (Fig 5A). In contrast, 94.8% (27,924) of genes assembled using the guided assembly approach showed perfect contiguity and completeness (Fig 5B). Increasing read length to 200 bases appeared to increase the number of genes with high *Cp* scores in the *de novo* approach (Fig 5C); however, only 14,252 genes (53%) in total had contiguity and completeness equal to 1. A much greater fraction of genes (98%; 27,861) with perfect contiguity and completeness were also observed for the guided assembly approach using reads of 200 bases (Fig 5D).

**Fig 5.**
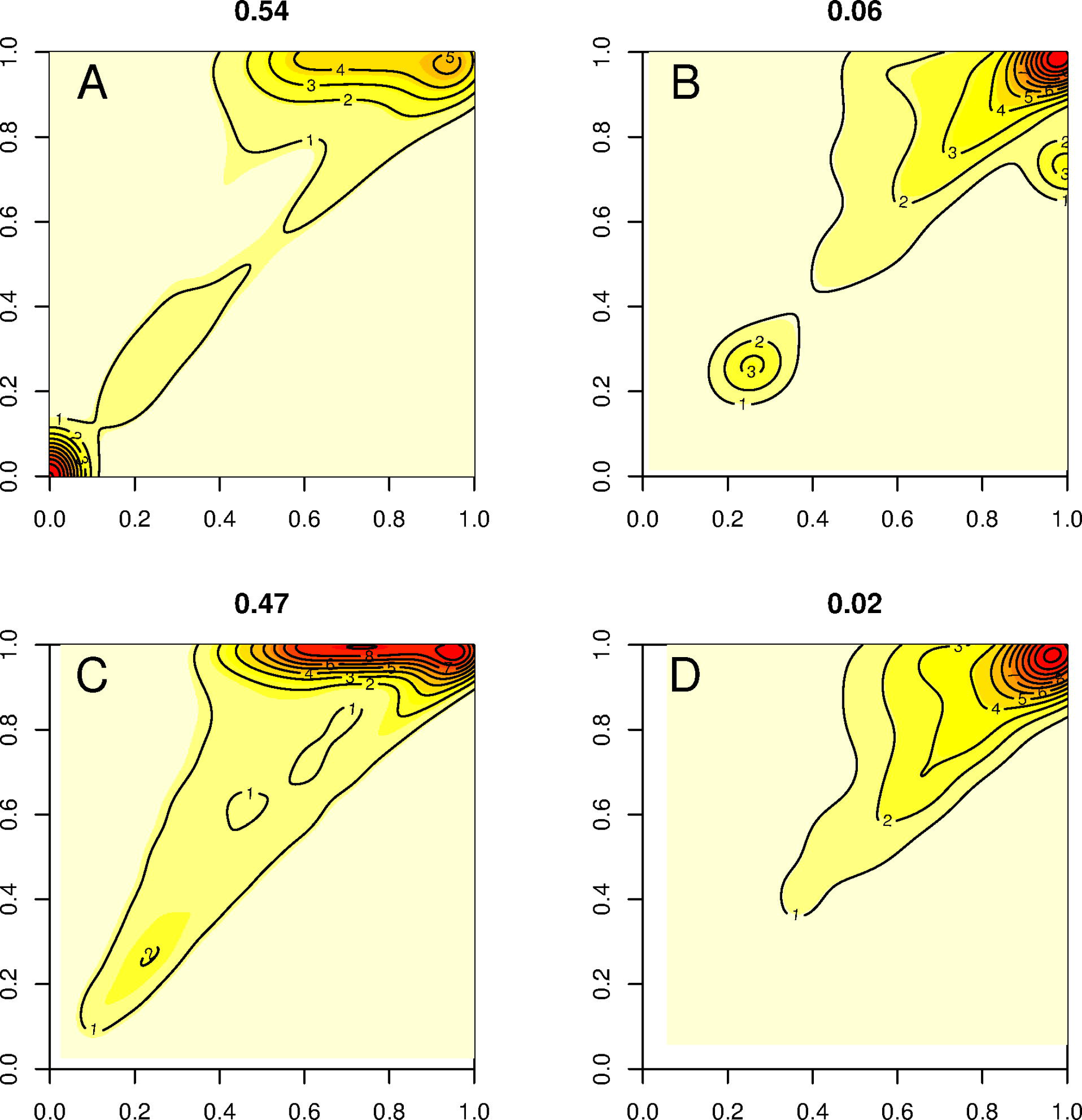
**Nonparametric estimation of gene density as a function of contiguity (x-axis) and completeness score (y-axis)** obtained from *de novo* and transcriptome-guided assembly pipelines for 0% divergence. Colors increasing from yellow to dark red denote increasing gene densities. Note also that for clarity of visualization, only the non-perfect fraction of genes are displayed. 100 base reads assembled *de novo* (A) and with guided assembly (B); 200 base reads *de novo* (C) and guided assembly (D). The proportion of non-perfect genes is indicated at the top of each panel.

Increasing divergence led to a drastic reduction in the number of genes with perfect *Ct* and *Cp* for both approaches, decreasing at 30% divergence to only 5.3% of *de novo* assembled genes and 5.7% of genes from the guided assembly approach using 100 base reads. However, approximately 80% of genes present contiguity and completeness above 0.95, for both methods (supporting information 7). Increasing read length did little to improve performance of either method, with only 5.5% of genes assembled from 200 base reads scoring perfect contiguity and completeness irrespective of pipeline used. Nevertheless, 85% of genes did present contiguity and completeness above 0.95 for both methods in this scenario (supporting information 7).

### Assembling non-model species transcriptomes

For *Parachondrostoma toxostoma*, a total of 18,519 genes were common to both methods. Nevertheless, we observed a clear difference in the number of detected genes between *de novo* and guided assemblies: a significantly higher proportion of genes were detected for *P. toxostoma* (*X^2^*=4.65, P<10^-5^) when using the guided assembly (20,605 genes) than for the *de novo* approach alone (20,032 genes). The performance of the guided assembly approach was also higher than *de novo* alone, as measured by both contiguity (supporting information 8A; t = -48.084, df = 80704, p-value < 2.2e-16) and completeness (supporting information 8A; t = -44.669, df = 80376, p-value < 2.2e-16). It should be noted that these metrics have a different meaning for empirical data than for simulated data: here they describe how the reconstructed contigs compared to the longest ones from *D. rerio* for a given gene.

For *Quescus pubescens*, a total of 8,886 genes were common to both methods. Nevertheless, a significantly higher proportion of genes were detected for *Q. pubescens* (*X^2^*=1873.808, P<10^-5^) when using the guided assembly (16,326 genes) than for the *de novo* approach alone (9,385 genes). The performances of the assembly approaches were quite similar as measured by contiguity (supporting information 8B; t = 4.2291, df = 18421, p-value = 2.357e-05), but did not differ significantly for completeness (supporting information 8B; t =1.5887, df = 18163, p-value = 0.1121). The efficiency of the guided assembly is high, particularly when compared with the *de novo* approach conducted by Torre et al. [26] who identified only 11,074 genes based on assignment to *V. vinifera*. These two transcriptome analyses underscore the high performance of our pipeline considering that 35.66% of the orthologous genes present a divergence from the reference transcriptome higher than 20% for *P. toxostoma* and 70.54% for *Q. pubescens* (supporting information 11). Moreover, there are some general trends in the two inferred transcriptomes. For example, the coverage does not depend on the length of the genes and is on average higher than 10x for all size classes (supporting information 12). Neither does completeness depend on the length of the gene (0.8 for *P. toxostoma* and 0.7 for *Q. pubescens*), although these values decrease slightly for gene lengths higher than 3,000 bases for *P. toxostoma* (25 % of genes). Converesely, we observed that coverage does impact the completeness when lower than 2.4X for *P. toxostoma* and 4.0X for *Q. pubescens*. Likewise, we observed that genetic divergence is negatively correlated with completeness.

## Discussion

### Read assignment with *blastn* and characterizing gene categories

We developed a pipeline based on the use of blastn to assign reads to reference transcriptome(s) and we tested this pipeline with simulated data over a range of read lengths and sequence divergence relative to the reference. Our analyses provide a much-needed empirical evaluation of the utility and limits of this approach, demonstrating how both read length and sequence divergence affect the efficiency and performance of read assignment. Irrespective of read length, and with 0% sequence divergence, 1,663 of the 31,944 simulated genes (5.21%) could not be retrieved. This result was surprising given that reads were simulated directly from the corresponding reference transcriptome, yet in the analysis, multiple assignments were retained as equal best hits. We observed that when a read equally matches two different genes, the BLAST score is highest for the gene that displays a length closest to the query read length, hence this gene acts as an attractor for those reads that may not be assigned to the correct gene.

High sequence divergence (30%) and short read lengths (100 and 150 bases) had a negative impact on recovery rate, substantially reducing the number of genes with a recovery rate *rr*=1 (i.e. all the generated reads being correctly assigned). The effect of sequence divergence is congruent with studies on *Drosophila* and primates [50], suggesting that 30% divergence may represent an upper threshold for the recovery of orthologous sequences. This is, however, much better than assignment methods based on mapping reads whose performance is capped at around 15% divergence [20]. When sequence divergence is non-null, increasing read length (200 and 350 bases) limits the erroneous assignment towards paralogous genes. Genes with perfect specificity (i.e. *sr*=1; no assigned read belongs to another *gene-id*) were less sensitive to either divergence or read length. Although few genes were classified as ‘recipient’ genes (*sr*<1), these could represent an important source of bias in quantitative analyses of RNA-seq data, appearing as over-expressed via the erroneous assignments from “donor” genes. It should also be noted that a recipient gene could act as an attractor for multiple donors, as evidenced by the high fraction of donors relative to recipients. Such genes would appear as highly over-expressed. Although these would obviously not lead to erroneous inference of over-expression in our pipeline (i.e. they would be removed/filtered prior to analyses), they could nevertheless contribute to an underestimation of the true levels of expression for donors. As such, we highly recommend identifying such problematic genes as a mean of distinguishing biologically meaningful signals from artefact in the interpretation of differential expression profiles from RNA-seq data. To this end, we have made all scripts used for simulating reads from a reference genome available on https://github.com/egeeamu/voskhod; these scripts can be modified for use on any other reference transcriptome used to annotate RNA-seq data, with read length and divergence parameters adjusted to match those under investigation.

Our analyses also highlight the difficulties that can arise when working with a reference transcriptome that is highly divergent from one’s focal, non-model species – this was facilitated by our *in silico* generation of reads that in turn enabled development of a matrix of gene categories (given the length of the transcripts fragments) based on recovery and specificity rates (parameters *rr* and *sr*). Specifically, we demonstrated that when sequence divergence increases to 30%, the proportion of ‘donor’ genes (*rr*<1, *sr*=1) and ‘mixed’ genes (*rr*<1, *sr*<1) increases, and the proportion of ‘perfect’ genes (*rr*=1, *sr*=1) decreases. Surprisingly, the proportion of ‘recipient’ genes (*rr*=1, *sr*<1) does not increase; however, their density within the genome likely does, particularly if analyses are based on short sequence reads. For example, when analyses are based on longer reads, recipient genes are largely derived from the smaller genes (i.e. <1,000 bases); however, short genes appear sensitive to the accumulation of recipients of various size (contrast Fig 3E and 3F). In this instance, gene length alone could not be used as a quality filter for excluding genes in downstream analyses of expression. Additionally, some genes previously classified as perfect become ‘donor’ genes at high divergence rate. Likewise, the percentage of ‘perfect’ genes decreased from 85.43% to 68.60% when read lengths where reduced from 350 to 100 bases. This result can be caused by i) a recently duplicated gene found at multiple loci in the genome and/or ii) a conserved domain (or repetitive domain) in gene families. Irrespective of the underlying causes, this result highlights the advantages in terms of increased confidence of assignment when working with longer reads, a point not to be neglected when planning RNA-seq experiments for non-model species.

### Improving transcriptome inference with transcriptome-guided assembly based on blastn

Our comparison of the efficiency and performance of *de novo* in combination with transcriptome-guided assembly based on blastn was based on three key metrics: the number of genes identified, contiguity and completeness. Although short reads yielded a higher number of genes identified, the quality of genes was lower for *de novo* alone with respect to contiguity and completeness (i.e. an increased proportion of segmented and/or fragmented transcripts). Moreover, the observed increase of these two metrics with increasing sequence divergence – a trend most pronounced in the *de novo* only approach – was certainly counter-intuitive, but explained by the fact that several *D. rerio* genes share identical sequences (paralogous genes and repetitive conserved domains). To some extent this might be expected in species that have undergone extensive genome duplication, such as cyprinids; the extent to which this trend is evident in species with more conserved genomes is a topic of potential future interest. Nevertheless, as we generated divergent sequences (up to 30%) with a random process, this artificially decreased the similarity between paralogous genes or conserved domains, and artificially enhanced assembly metrics. At 0% divergence, the efficiency (i.e. percentage of genes identified) of *de novo* assembly is less impacted than its performance (contiguity and completeness). Several strategies are available to optimize d*e novo* assembly, for example using multiple K-mer values in a de Bruijn graph to handle both over- and under-expressed transcripts [37,43]. Evaluating various optimization strategies is beyond the scope of this article, but as we propose a solution using an assembly based on read assignment, an improvement in any one of the constituent parts of the pipeline would benefit the entire transcriptome inference.

As a guideline, we would recommend using the combination of *de novo* and transcriptome-guided assembly based on blastn, as this systematically increases the number of identified genes relative to *de novo* alone, although there was a slight decrease at 30% divergence. Even when query and reference are identical (i.e. 0% divergence), the guided assembly approach yields advantages, with 98% of identified genes displaying contiguity and completeness equal to 1. Moreover, a higher percentage of genes are successfully retrieved using the guided-assembly approach implemented here (88.1%) than for the *de novo* assembly approach alone (81.5%). As far as performance is concerned, the guided-assembly outperforms *de novo* assembly alone for both contiguity and completeness. When divergence increases towards 30%, both approaches converge to the same results. This overall greater efficiency and performance of the transcriptome-guided assembly based on blastn allows identifying more genes with a higher degree of confidence in associated assignments. Combining a *de novo* assembly and a related reference transcriptome for read assignment also addresses the bias/error in contigs caused by the dependence on a related reference alone. Empirical data corroborate these findings for both non-model species analyzed here. When assembling the *Parachondrostoma toxostoma* transcriptome, of the 31,944 genes known from *D. rerio*, the guided and *de novo* assemblies recover respectively 20,605 and 20,032 genes, but the performance of the guided assembly approach is much higher for both the contiguity and completeness metrics. For *Q. pubescens*, the performance was similar for the two assembly approaches, but the efficiency of the new combined method clearly outperformed *de novo* alone with almost twice the number of genes detected.

Altogether, with the categorization of genes relative to their assignment behavior and its consequences on sorting relevant genes for further analyses, combining a *de novo* step with the use of blastn to assign reads prior to a guided assembly significantly improves the quality of the reconstructed transcriptomes for non-model organisms.

## Acknowledgments

We specially thank Bernard Barascud for his great involvement with the common garden experiments.

## Supporting information captions

S1 Table: output example of the transcriptome-guided assembly pipeline (simulations) corresponding to a collection of database entries displaying the number of identified reads for each gene-id.

S2 Protocol: Simulating reads for efficiency and performance testing.

S3 Text: Statistical framework for evaluating reads assignment performance.

S4 Protocol: Illumina library production.

S5 Table: Prediction of true assignment probability for recovery rate and specificity rate.

S6 Table: Gene simulation of reads assignment to the *D. rerio* transcriptome using Blastn and showing the five different categories (mixed, donor, recipient, perfect, undetectable).

S7 Figure: Nonparametric estimation of the contiguity (x-axis) and completeness score (y-axis).

S8 Figure: Boxplots of contiguity and completeness.

S9 Figure: Biological processes of the identified genes.

S10 Text: Computing resources and computation time.

S11 Figure: Genetic divergence between genes from reference and non-model organisms.

S12 Figure: Interaction between coverage, gene size classes and completeness for *Parachondrostoma toxostoma* and *Quercus pubescens*.

